# Agent-Based Modeling Predicts HDL-independent Pathway of Removal of Excess Surface Lipids from Very Low Density Lipoprotein

**DOI:** 10.1101/398164

**Authors:** Yared Paalvast, Jan Albert Kuivenhoven, Barbara M. Bakker, Albert .K. Groen

**Affiliations:** Laboratory of Pediatrics, University of Groningen, University Medical Center Groningen, 9713 AV Groningen, The Netherlands; Department of Laboratory Medicine, University of Groningen, University Medical Center Groningen, 9713 AV Groningen, The Netherlands; Laboratory of Experimental Vascular Medicine, University of Amsterdam, Amsterdam UMC, location Meibergdreef, 1105 AZ Amsterdam, The Netherlands

## Abstract

A hallmark of the metabolic syndrome is low HDL-cholesterol coupled with high plasma triglycerides (TG), but it is unclear what drives this close association. Plasma triglycerides and HDL cholesterol are thought to communicate through two distinct mechanisms. Firstly, excess surface lipids from VLDL released during lipolysis are transferred to HDL, thereby contributing to HDL directly but also indirectly through providing substrate for LCAT. Secondly, high plasma TG increases clearance of HDL through core-lipid exchange between VLDL and HDL via CETP and subsequent hydrolysis of the TG in HDL, resulting in smaller HDL and thus increased clearance rates.

To test our understanding of how high plasma TG induces low HDL-cholesterol, making use of established knowledge, we developed a comprehensive agent-based model of lipoprotein metabolism which was validated using monogenic disorders of lipoprotein metabolism.

By perturbing plasma TG in the model, we tested whether the current theoretical framework reproduces experimental findings. Interestingly, while increasing plasma TG through simulating decreased lipolysis of VLDL resulted in the expected decrease in HDL cholesterol, perturbing plasma TG through simulating increased VLDL production rates did not result in the expected HDL-TG relation at physiological lipid fluxes. However, model perturbations and experimental findings can be reconciled if we assume a pathway removing excess surface-lipid from VLDL that does not contribute to HDL cholesterol ester production through LCAT. In conclusion, our model simulations suggest that excess surface lipid from VLDL is cleared in part independently from HDL.

**Author summary:** While it has long been known that high plasma triglycerides are associated with low HDL cholesterol, the reason for this association has remained unclear. One of the proposed mechanisms is that during catabolism of VLDL, lipoproteins rich in triglyceride, the excess surface of these particles become a source for the production of HDL cholesterol, and that therefore decreased catabolism of VLDL will lead to both higher plasma triglyceride and low HDL cholesterol. Another proposed mechanism is that during increased production of VLDL, there will be increased exchange of core lipids between VLDL and HDL, with subsequent hydrolysis of the triglyceride in HDL, leading to smaller HDL that is cleared more rapidly. To investigate these mechanisms further we developed a computational model based on established knowledge concerning lipoprotein metabolism and validated the model with known findings in monogenetic disorders. Upon perturbing the plasma triglycerides within the model by increasing the VLDL production rate, we unexpectedly found an increase in both triglyceride and HDL cholesterol. However, upon assuming that less excess surface lipid is available to HDL, HDL decreases in response to increased VLDL production. We therefore propose that there must be a pathway removing excess surface lipids that is independent from HDL.

**Abbreviations:**

PR: (production rate)
FCR: (fractional catabolic rate)
ppd: (pool per day)
SRB1: (scavenger receptor B1)
EL: (endothelial lipase)
HL: (hepatic lipase)
PLTP: (phospholipid transfer protein)
CETP: (cholesteryl ester transfer protein)
FC: (free cholesterol)
CE: (cholesterol ester)
PL: (phospholipid)
LpX: (lipoprotein X).

## Introduction

High plasma triglycerides (TG) and low HDL-cholesterol (HDL-C) are important risk factors for cardiovascular disease [1,2]. Which of these two strongly negatively correlated parameters confers cardiovascular disease risk is still a matter of controversy [3]. It has been suggested that lipolytic activity mediates the correlation between HDL-C and plasma TG [4]. According to this hypothesis triacylglycerol lipolytic activity increases HDL-C as it allows apolipoproteins, phospholipids (PL) and free cholesterol (FC) to be transferred to the HDL-pool [5]. An alternative explanation holds that in conditions with high plasma TG, cholesterol ester transfer protein (CETP) – mediated exchange of HDL – cholesterol ester (CE) with TG from TG-rich lipoproteins is enhanced, thereby decreasing HDL-C [6]. Which of these mechanisms is more important and under what conditions is unclear. The HDL-C and TG plasma concentrations measured are the end result of a multitude of interacting processes. A short introduction to metabolism of apoA-I and apoB-containing lipoproteins is given below.

### HDL metabolism

The production of HDL not only depends on the production of its major protein, apolipoprotein (apo) A-I, but also on the activity of ATP binding cassette subfamily A member 1 (ABCA1), and lecithin-cholesterol acyltransferase (LCAT) [7]. Maturation of lipid-poor apoA-I to larger HDL is dependent on LCAT activity on the particle’s surface, resulting in the change of a discoidal 2 or 3 apoA-I molecules harboring disc with PL and FC, to a more spherical particle containing CE in its core [8]. These matured spherical HDL-particles are subject to fusion, a process catalyzed by phospholipid transfer protein (PLTP). This results in both larger particles containing typically 3 to 5 apoA-I molecules as well as shedding of superfluous apoA-I as lipid-poor apoA-I [9]. ApoA-I shedded in this way can then be either recycled for formation of new spherical HDL, or be cleared by the kidneys [10]. Furthermore, HDL interacts with apoB-lipoproteins through CETP-mediated exchange of TG by CE [11]. TG in the core of HDL can then be hydrolyzed by both hepatic (HL) and endothelial lipase (EL) [12,13]. Depletion of the HDL-core will lead to further shedding of lipid-poor apoA-I, which, as stated already, can then either be cleared by the kidneys or be remodeled back into mature HDL [14]. The specific uptake of HDL-CE by the liver through scavenger receptor class B member 1 (SRB1) followed by excretion into the bile is considered to be an important mechanism for cholesterol export from the body. The existence of all these pathways has been demonstrated almost exclusively in *in vitro* systems. Whether they are important under *in vivo* conditions has seldom been shown and whether lipoprotein fluxes are in balance when these pathways exclusively describe lipoprotein metabolism has never been shown.

### Metabolism of triglyceride-rich lipoproteins

TG-rich lipoproteins, which may be either very-low density lipoprotein (VLDL) produced by the liver or chylomicrons produced by the small intestine, have their triglycerides hydrolyzed by lipoprotein lipase (LPL) in peripheral tissues, becoming progressively smaller as this process continues. For VLDL, HL is considered to facilitate the removal of the remaining portion of TG at the end of this lipolytic cascade [15]. The resulting LDL is mainly taken up in the liver through the action of LDL-receptor (LDLR), thus the same tissue in which apo-B containing lipoproteins are originally produced. In addition to LDLR-mediated uptake, there is also some uptake through LDLR-related protein (LRP) and VLDL-receptor (VLDLR), that facilitate uptake of larger apoB-lipoproteins that have not completely gone through the lipolytic cascade[16].

### Models of lipoprotein metabolism

To get a grip on the complexity of lipoprotein metabolism, several mathematical models have been developed. For example, Wattis et al. explored the changes in VLDL and LDL-size in relation to uptake by the LDL-receptor [17]. Van de Pas et al. created a model that describes whole body cholesterol homeostasis for both man and mouse [18]. In contrast, Schalkwijk et al. focused on the lipolytic cascade of only apoB-lipoproteins [19]. More recently, Lu et al. reported a mathematical model focusing on the interaction between apoB and apoA-I carrying lipoproteins and included remodeling of HDL [20]. All these models made use of ordinary differential equations (ODE). This type of models are well-suited to describe the kinetics of well-mixed pools of well-defined metabolites. Lipoproteins, however, are not a well-defined set of metabolites, but rather heterogenic bundles containing multiple protein and lipid species that continuously change in composition and size. In ODE-models this issue is usually circumvented by defining a broad range of lipoprotein classes, where each class represents a homogeneous pool with a pre-defined composition, so that for all practical purposes the lipoproteins can then be treated as if they were metabolites in a metabolic pathway. Such a strategy, however, limits the possibilities of studying the lipid exchange between actual lipoproteins. Interestingly, Hübner et al. explored the use of stochastic equations and the Gillespie algorithm to model how lipid-exchange between individual lipoprotein complexes may alter the lipoprotein profile, and may be regarded as a first effort in applying agent-based concepts on models of lipoprotein metabolism [21]. Such an agent-based approach does not require pre defining of lipid compositions for a specific lipoprotein class, but allows for creating a large number of ‘lipoprotein agents’ that can each have a different composition and interact with one another independent of the other agents. By giving the agents interaction rules based on current biological insights, new insights may be gained by observing how these rules affect the system as a whole [22].

The aim of this study is to test existing hypotheses that explain the inverse correlation between TG and HDL-C. To this end, we created an agent-based model and validated the model with findings from patients with monogenetic disorders in lipoprotein metabolism. The validated model was subsequently used to test existing hypotheses concerning the association between plasma TG and HDL-C. It was first of all found that the presence of CETP-activity alone is not sufficient to explain this relationship. Furthermore, we found that we had to drop the assumption that all surface lipids released during lipolysis of triglyceride-rich lipoproteins are contributing to HDL-CE in order to explain results obtained through lipoprotein kinetic experiments, showing that increased VLDL-production rate (PR) is associated with increased apoA-I fractional catabolic rate (FCR). We discuss how these findings fit in the established but largely theoretical framework of lipoprotein metabolism and conclude that a pathway must exist that clears surface lipids from VLDL in a manner that is independent of HDL.

## Methods

In the constructed agent-based model, lipoproteins are represented as patches on a grid, where properties of those patches can be changed independently of one another. In this way, one-on-one interactions between heterogenic particles become possible and thus accessible for study. The model contains all the elements considered important for lipoprotein metabolism (Fig 1). A full description may be found in the appendix (Suppl. File S1), and the code of the model may be examined by opening the model in NETLOGO [22] and clicking on the ‘code’-tab (Suppl. File S2). We briefly discuss the main points here.

**Fig 1.**
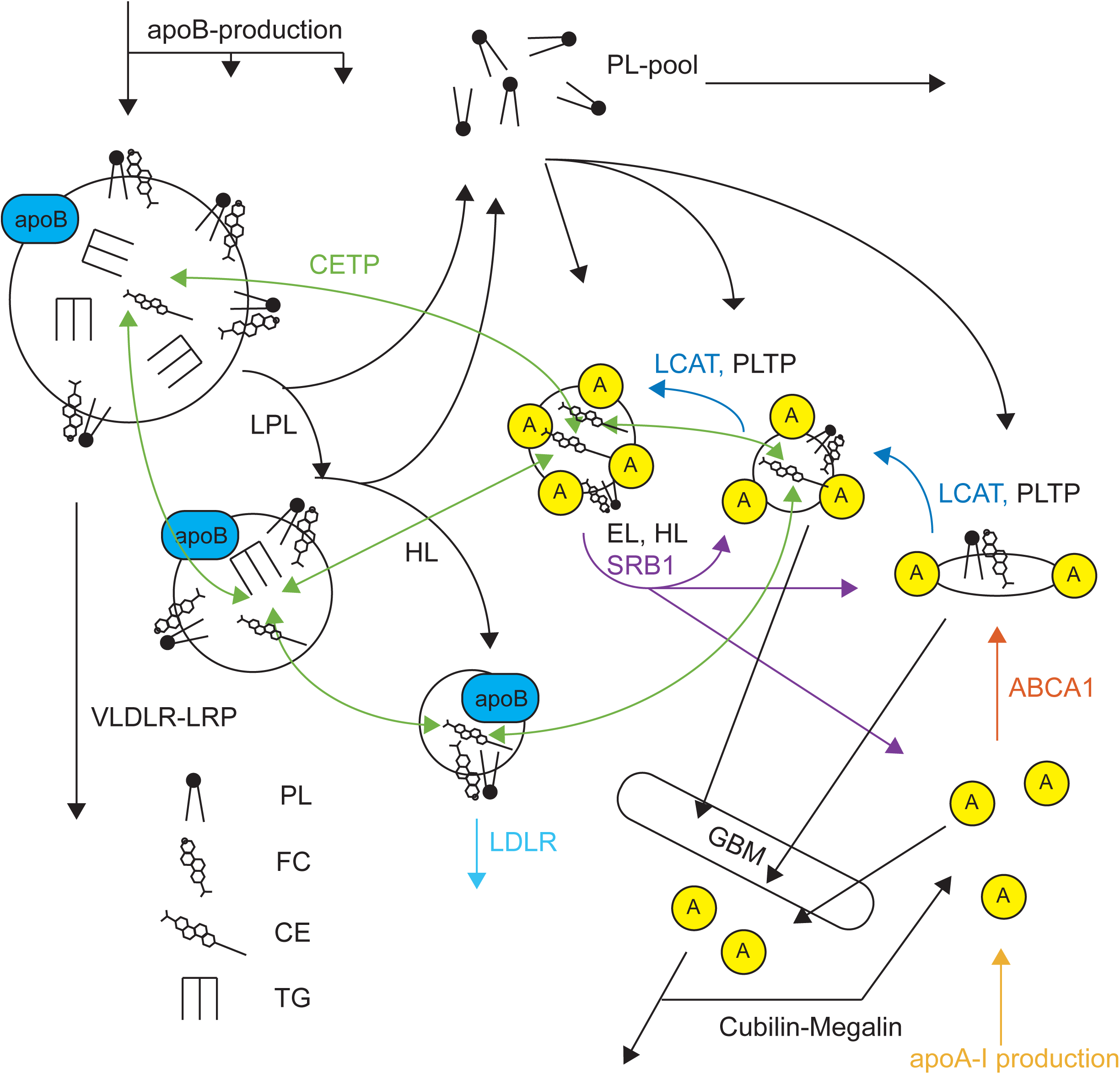
Overview of the model structure.

ApoB-carrying lipoproteins enter the model as VLDL, and after lipolysis by LPL and HL, are taken up by LDLR. During lipolysis, these lipoproteins become smaller and shed phospholipid. This phospholipid is redistributed to the HDL-compartment, where it can function as a substrate for LCAT to form cholesterol ester and create larger HDL, which is cleared more slowly than small HDL due to the size-selectivity of the glomerular basement membrane (GBM) in the kidney. Not all phospholipids end up in HDL, however, and our agent-based model indicates that a significant portion is cleared through another route. Throughout, all lipoproteins are subject to remodeling by core lipid exchange through CETP, and HDL is remodeled through removal of PL and TG by EL and HL, and removal of core-lipids by SRB1. Such remodeling can lead to shedding of lipid-poor apoA-I that can be either be used for production of HDL, or be cleared by the GBM. The model components LCAT (blue), ABCA1 (orange), apoA-I (yellow), SRB1 (purple), CETP (green) and LDLR (light-blue) were used for validation.

### Lipoproteins represented by patches on a grid

In the NETLOGO programming language any actions in the model simulation take place on a grid with a pre-defined size. This grid consists of squares that can be accessed through their respective x-y coordinates and are referred to as patches. Patches as a class can have different attributes, and every patch is an instance of this class. The model assigns properties to patches (such as number of CE, or TG molecules) so that they may be treated as virtual lipoproteins. Because modeling individual lipoproteins is computationally infeasible, each patch (in silico) represents a bundle of lipoproteins with the same properties in vivo. Within the model, these patches are sorted for lipoprotein size along the x-axis, so that something akin to a lipoprotein profile is obtained, and a visual representation of what is happening in the model is obtained in ‘real-time’. It must be noted that during simulation not all patches represent lipoprotein bundles, and in fact many are ‘empty’. Furthermore, since patches themselves are defined by their x-y coordinates, they do not actually move over the grid during simulation, rather their properties are copied to an empty patch should the model logic dictate the simulation to do so. Finally, for these patches to interact with one another they do not necessarily have to be adjacent, and interaction between patches is fully controlled by the various algorithms and the stochastic processes within.

### V(LDL) production, composition and size

The model includes production of apoB-lipoproteins in the form of VLDL, IDL, and LDL. The production rate of these is based on findings in tracer-kinetic studies [23]. For calculation of the composition of the lipoproteins spherical shapes were assumed and a CE content of nascent VLDL half that of what is measured in LDL [24,25]. Every VLDL contains FC and PL as surface lipids and TG and CE as core lipids, so that at hepatic secretion VLDL has a size of 54 nm. Within the model, the rate of VLDL secretion (VLDL-PR) and the VLDL size can be changed independently by modulating the rate at which apoB patches are produced and by changing the TG content of VLDL respectively. The clearance of apoB-lipoproteins is mediated in part through the VLDLR-LRP – parameter, and was modeled to clear lipoproteins with a size greater than 32 nm in diameter. The major pathway of apoB clearance however is through the LDLR pathway, which was modeled so that it has a Gaussian distribution of affinity with relation to size, with optimal absorption of apoB lipoproteins with a size of 21.5 nm.

### Hydrolysis of triglycerides and phospholipids

TG is hydrolyzed from the core of the apoB-carrying lipoproteins by LPL, upon which they will shed excess surface lipids. The activity of LPL in the model depends on the number of apoB lipoproteins associated with the enzyme; the chance to associate with LPL is proportional to the surface area, and the chance to dissociate from LPL increases once the particle obtains a size in the range of IDL (25 nm). HL-activity also depends on the number of lipoproteins associated with HL but the chance of association with HL was modeled to be normally distributed around the size of 32 nm. The chance that HL dissociates from the lipoprotein increases upon reaching LDL-size (21.5 nm). During the lipolytic cascade, excess surface is funneled into an intermediate PL-pool from where the PL is redistributed to HDL, i.e. apoA-I carrying lipoproteins. Based on the literature, it is assumed that approximately 65% of the excess surface lipid from VLDL is available to HDL, which is a conservative estimate [26,27].

### Production, remodeling and clearance of HDL

ApoA-I enters the system as lipid-poor apoA-I, which interacts with ABCA1 to form nascent HDL-particles that carry 2 to 3 apoA-I molecules and in the order of 180 phospholipid molecules as well as 130 free cholesterol molecules, and virtually no core lipids [28]. These nascent HDL particles can then be converted to more spherical and larger HDL by action of LCAT, which transfers a fatty acid chain from PL to FC on the surface to form CE. Activity of LCAT is constrained by the amount of surface lipid because the model does not allow for less surface lipid than would be required to fully envelop the volume of core lipids. Clearance of core lipids occurs through activity of SRB1, reducing the size, and producing a chance to shed some lipid-poor apoA-I. The clearance of apoA-I on the other hand, occurs through the filtering of lipid-poor apoA-I and HDL through the glomerular basement membrane in the kidneys, which was modeled as a function of glomerular filtration rate and size of the lipoprotein, so that lipid-poor apoA-I is very likely to be filtrated, whereas large HDL is very unlikely to be filtrated [29]. Once filtrated, the cubulin-megalin parameter controls the chance of the filtrated apoA-I to be reabsorbed and recycled, thus negating the clearance and reintroducing the filtered HDL as an equivalent number of apoA-I [10]. Other than by SRB1 and LCAT, HDL in the model is further subject to remodeling by HL and EL, which remove mainly TG and PL from the particle respectively. Moreover, there is activity of PLTP, which facilitates the fusion of two HDLs, creating a larger HDL with more surface, that can then also shed some of its superfluous lipid-poor apoA-I. The shedding of lipid-poor apoA-I given a certain core volume, and the amount of phospholipid HDL may carry (given a number of apoA-I), was based on a mathematical model developed by Shen, Mazer and colleagues and colloquially referred to as ‘’Shen’s model’’ [20,30].

### CETP

Finally, lipoproteins are remodeled through activity of CETP, that facilitates the exchange of TG and CE between all classes of lipoproteins. Algorithms considered for our model focused on the chemical potential as driving force (*Alg1*) [31], and a more phenomenological algorithm where TG and CE transfer were uncoupled and a net transfer of CE from HDL to V(LDL), and a net transfer of TG from V(LDL) to HDL was assumed (*Alg2*) [32]. The algorithm (*Alg3*) used in the final model amalgamated these two ideas by using both chemical potential and hydrostatic pressure as the driving force between exchange of core lipids between lipoproteins [33].

### Stochasticity, iterations and run-time

Since the model contains stochastic elements, every simulation run is repeated with five different seeds for the random number generator. Unless stated otherwise, only the parameter under study is varied and for other parameters the default values are used.

## Results and Discussion

### Default parameters in the model results in states and fluxes within the physiological range

Using the default parameters for the model and allowing the system to reach steady state by simulating a time course of three weeks, allowed for evaluating whether the parameters lead to states and fluxes that are within a range that would be expected under physiological conditions. Simulating a trajectory of three weeks led to a pseudo-steady state under all the tested conditions (Fig. 2). Parameters were varied manually to find an appropriate set of default parameters. At default parameters, we find steady state values for both states and fluxes that are in reasonable agreement to values measured *in vivo* (Table 1).

**Fig 2.**
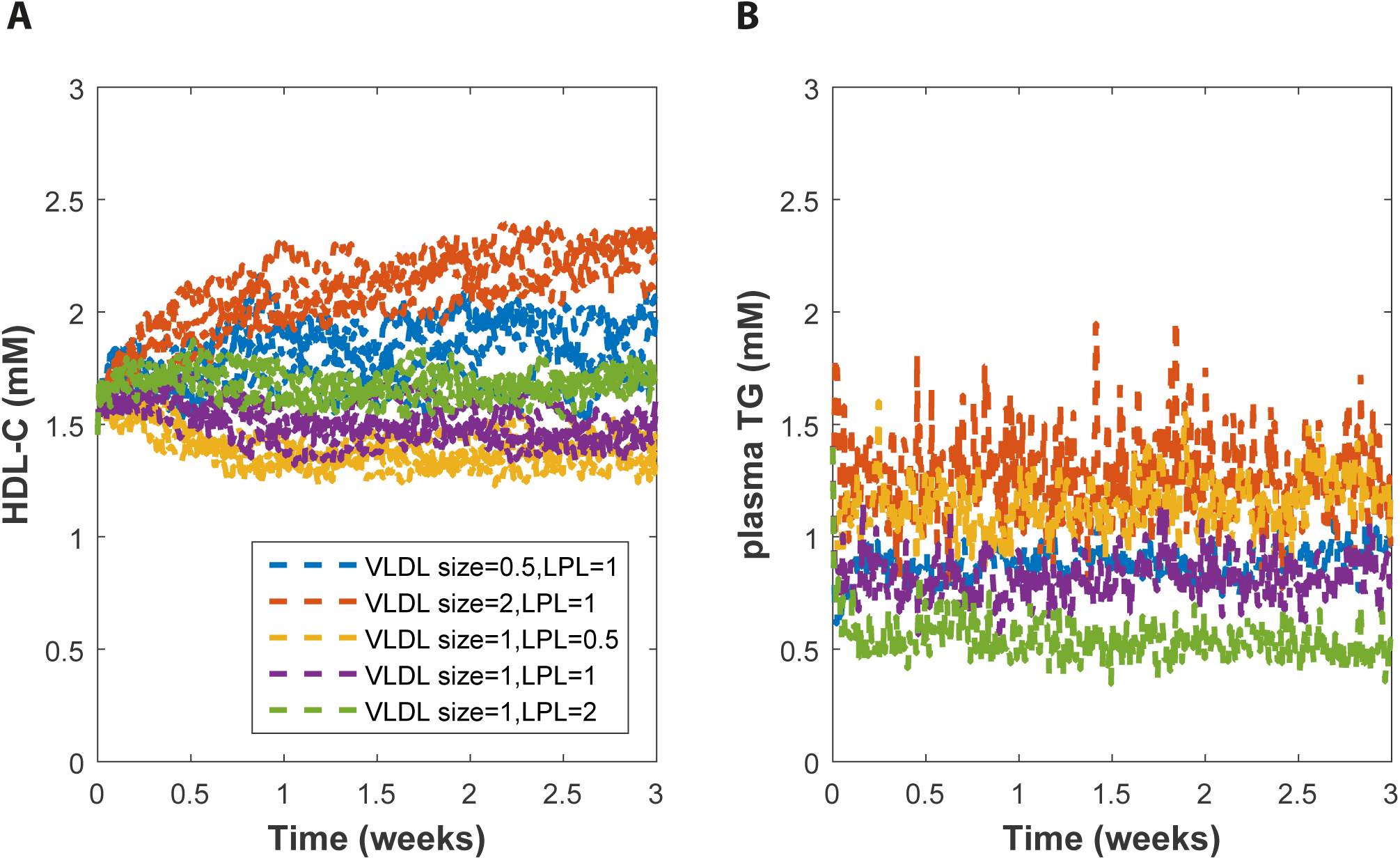
Simulation of a three weeks’ time course.

Simulation of a three weeks’ time course for different parameter values for VLDL size and activity of LPL, where VLDL size = 1 and LPL = 1 are default parameter settings. HDL-C is shown for 5 runs for each condition, whereas for plasma TG only the average of 5 runs is shown because the higher variability would make it the figure illegible. Note that plasma TG reaches steady state almost instantaneous whereas HDL-C, in agreement with a turnover time of approximately 5 days, takes much longer.

**Table 1.**
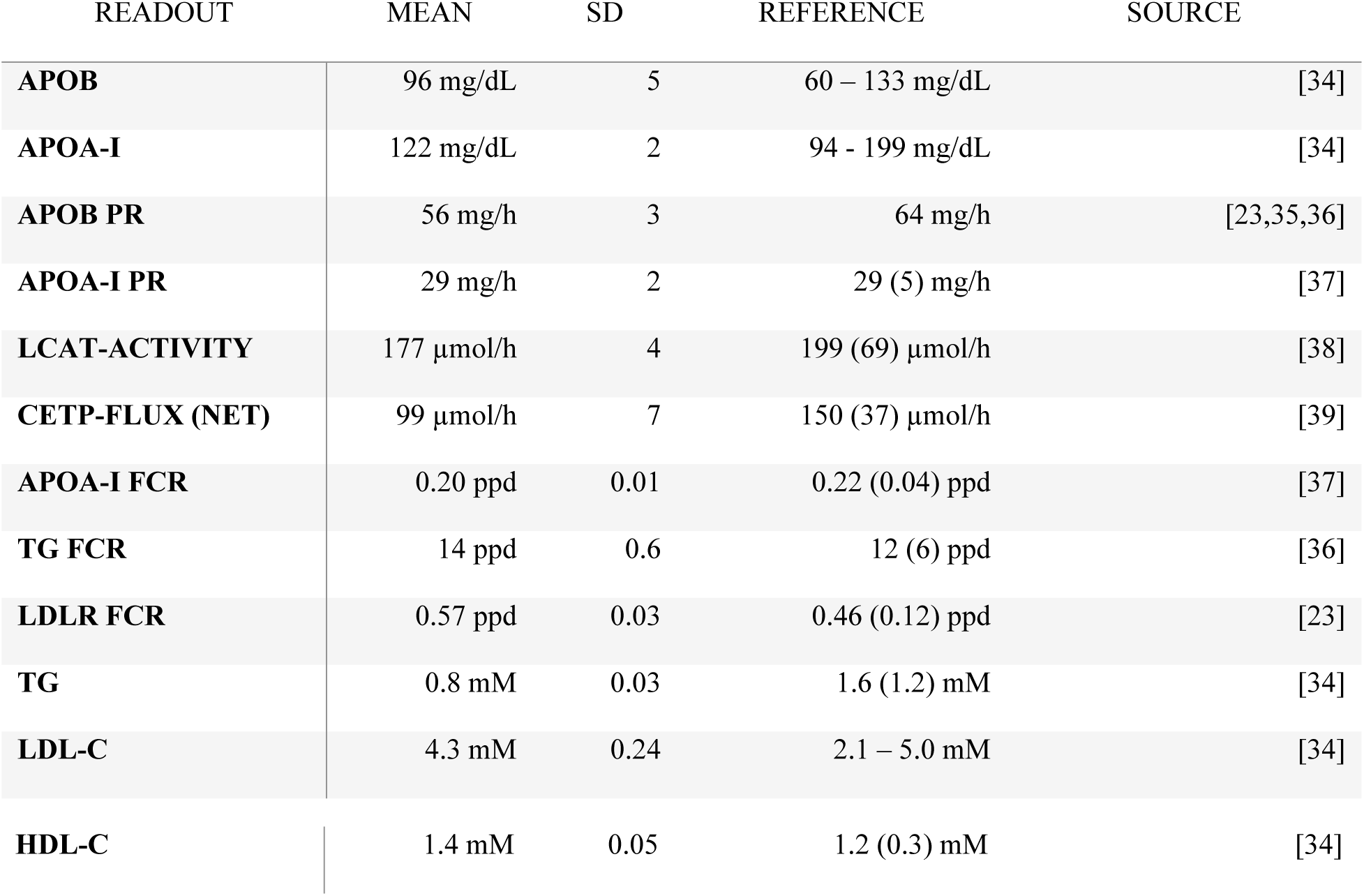
Steady state values at default parameters in the model and concomitant reference values.

### Model validation with patient data

To validate the model, we tested the effect of heterozygosity or homozygosity/compound heterozygosity for loss of function mutations in major lipid genes (*LCAT, ABCA1, APOA1, SCARB1, CETP, LDLR*) by setting the respective parameter at 50% or 0%, respectively. When varying one of these parameters, all other parameters were kept at their default value. The model simulations were in the majority of cases close to findings in affected individuals (Table 2 and Fig. 3), demonstrating that the model allows for studying lipoprotein metabolism over a wide range of conditions.

**Fig 3.**
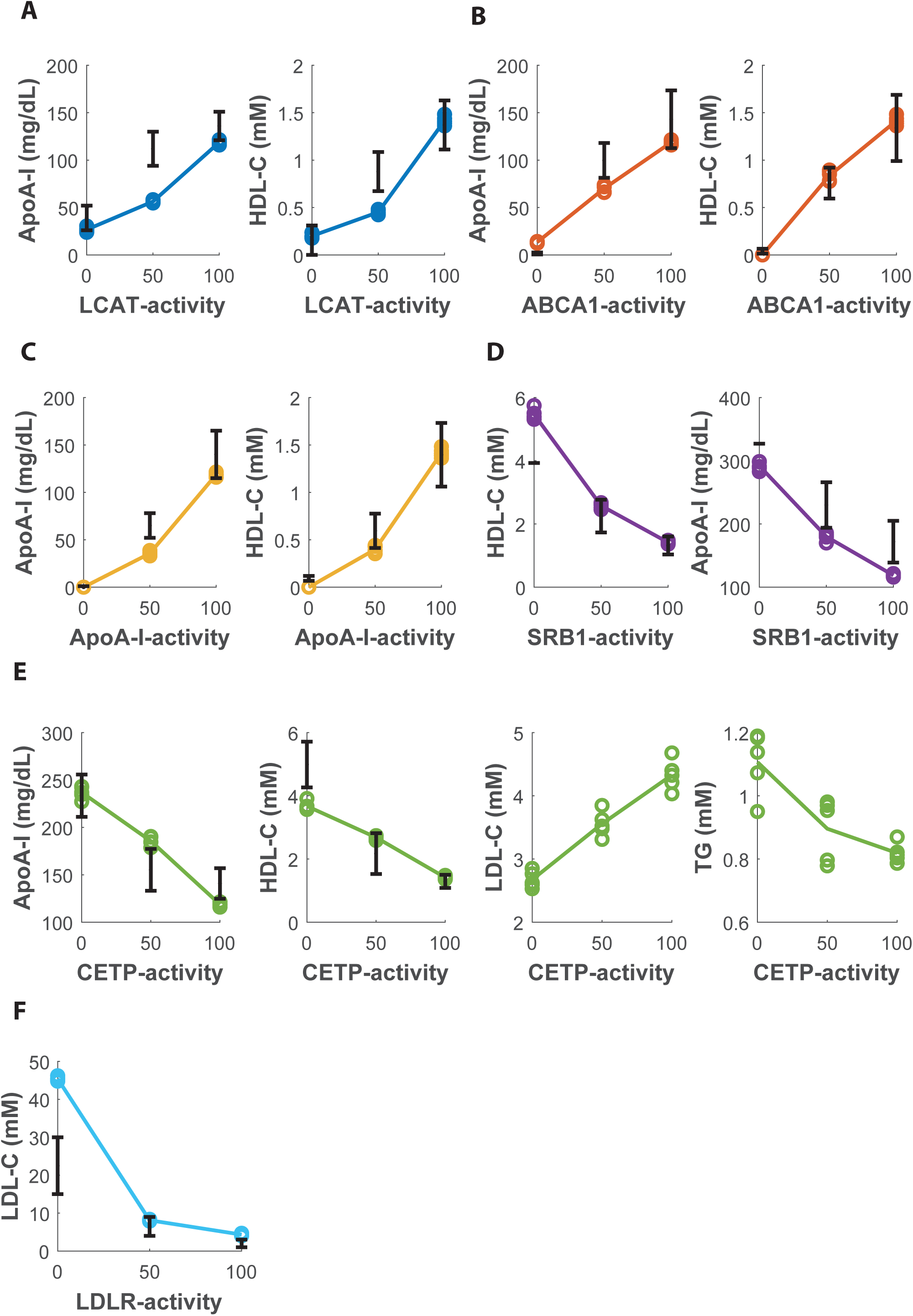
Simulating for loss-of-function mutations in genes important for lipoprotein metabolism.

Effect of varying LCAT, ABCA1, apoA-I, SRB1, CETP and LDLR-activity. Activity of 100% reflects controls, 50% of heterozygotes for loss of function mutations, and 0% for patients with complete deficiencies (homozygosity or compound heterozygosity for loss of function mutations) in the respective protein. The error bars reflect actual findings in controls, heterozygotes and homozygotes for the defects in the respective genes. Colored circles and lines reflect individual model simulations and means of those simulations, respectively. Note how all model simulations are within reasonable range of the actual findings in humans. No error bar is depicted for SRB1-deficiency because only one patient has so far been reported.

**Table 2.** Monogenetic disorders and model simulation results.

| GENE | READOUT | HOMOZYGOUS<br>(PATIENTS) | HOMOZYGOUS<br>(MODEL) | HETEROZYGOUS<br>(PATIENTS) | HETEROZYGOUS<br>(MODEL) | CONTROLS<br>(SUBJECTS) | CONTROLS<br>(MODEL) | REFERENCE |
| --- | --- | --- | --- | --- | --- | --- | --- | --- |
| **LCAT** | HDL-C (mM) | 0.05-0.3 | 0.3 (0.02) | 0.88 (0.2) | 0.5 (0.02) | 1.3 (0.3) | 1.4 (0.05) | [40–42] |
| **LCAT** | apoA-I (mg/dL) | 26-52 | 30 (2.4) | 112 (18) | 56 (1.5) | 136 (15) | 119 (2) | [40–42] |
| **APOAI** | HDL-C (mM) | 0.1 (0.02) | 0 (0) | 0.6 (0.2) | 0.5 (0.04) | 1.4 (0.3) | 1.4 (0.05) | [43] |
| **APOAI** | apoA-I (mg/dL) | 0 | 0 (0) | 65 (13) | 40 (2.2) | 140 (25) | 119 (2) | [43] |
| **ABCA1** | HDL-C (mM) | 0.04 (0.03) | 0 (0) | 0.7 (0.2) | 0.6 (0.05) | 1.2 (0.4) | 1.4 (0.05) | [44] |
| **ABCA1** | apoA-I (mg/dL) | 1.5 (1) | 10 (0.8) | 95 (17.3) | 60 (3.7) | 143.3 (32.4) | 119 (2) | [44] |
| **SRB1** | HDL-C (mM) | 3.9 | 2.4 (0.17) | 2.3 (0.5) | 1.8 (0.09) | 1.3 (0.3) | 1.4 (0.05) | [45] |
| **SRB1** | apoA-I (mg/dL) | 327 | 200 (6.7) | 230 (36) | 158 (6.1) | 172 (33) | 119 (2) | [45] |
| **CETP** | HDL-C (mM) | 5.0 (0.7) | 4.6 (0.14) | 2.2 (0.6) | 2.1 (0.06) | 1.3 (0.2) | 1.4 (0.05) | [46] |
| **CETP** | apoA-I (mg/dL) | 233.5 (22.3) | 259 (6.5) | 155.3 (22.1) | 184 (4.8) | 140.9 (16.1) | 119 (2) | [46] |
| **LDLR** | LDL-C (mM) | 15-30 | 45 (0.5) | 4-9 | 8.1 (0.2) | 2.1-5.0 | 4.3 (0.2) | [47–49] |

Simulations of the homozygous LCAT loss-of-function mutations matches up with *in vivo* findings in patients, but model simulations for the heterozygous case underestimate apoA-I and HDL-C compared with human data [40–42]. Simulations for both apoA-I and ABCA1-deficiency, the other two essential elements in HDL-production, show full agreement for both homozygous and heterozygous cases [43,44]. On the clearance side of HDL, we find that simulations for SRB1-mutation show agreement with HDL-C and apoA-I for the heterozygous cases. The single homozygous patient has been described [45] to date makes it difficult to validate our predictions. ApoA-I values for SRB1 simulations appear to be higher overall in patients compared to simulations, however, we note that apoA-I measurements in the study on the loss-of-function SRB1-mutation were consistently higher than in other studies, since controls showed apoA-I values of approximately 170 mg/dL compared to 140 mg/dL in others (Table 2). Simulating CETP-deficiency leads to HDL-C and apoA-I levels in line with human data [46], and a decrease in LDL-C reminiscent of the effect of CETP-inhibitors [50]. Finally, the simulations for LDLR are well in line for the heterozygous cases, but LDL-C is overestimated in our simulations for homozygous cases [47–49]. Apparently some adaptive processes take place *in vivo* that are not fully accounted for in our model.

### Sensitivity analysis

Exploring the sensitivity of TG, and HDL-C for the various parameters, we varied the parameters between 0 and 10 times the default value. As may be expected, we then find that TG is mostly sensitive to VLDL-PR, VLDL-size, LPL, and to a lesser extent also for LDLR, CETP and HL (Fig. S2), while HDL-C is mostly sensitive to apoA-I production, LCAT, ABCA1, ABCA1x, SRB1, CETP, EL, cubulin-megalin, and the glomerular filtration rate (Fig. S3). Because HDL-C and TG in the model communicate through CETP and PL-transfer, we also observed parameters that mainly control TG, like VLDL-PR and LPL, to have an effect on HDL-C, and vice versa, parameters important for HDL-C, such as LCAT and SRB1, to affect steady state plasma TG values.

## The HDL – TG correlation

### Testing the inverse correlation between HDL-C and TG

Having established that the model is capable of reproducing findings in patients with mutations in six genes involved in lipoprotein metabolism, we used it to study the relation between HDL-C and plasma TG. The relation between these parameters has been thoroughly studied in metabolic syndrome patients that were administered stable isotope-enriched leucine and glycerol to trace kinetics of apolipoproteins and TG. These studies found strong correlations between increased plasma TG and increased VLDL-PR and relatively weak correlations with decreased clearance of VLDL [51,52]. It has furthermore been reported that production of large VLDL, specifically VLDL1, is increased in patients with metabolic syndrome [51]. Moreover, increased VLDL-PR is said to decrease HDL-C through increasing apoA-I FCR [51–53]. Finally, it is well-established that a decrease in LPL-activity also leads to an increase in plasma TG and a concomitant decrease in HDL-C [54–56].

Based on these tracer-kinetic studies, we expect that modulating plasma TG through decreasing LPL-activity will lead to a negative correlation between plasma TG and HDL-C in our model. We furthermore expect that increasing VLDL-PR or VLDL-size, will lead to higher plasma TG, and lower HDL-C. Now, by testing the model against these expectations, we can learn whether our model, and therefore our understanding of the lipoprotein metabolism, is sufficient to explain the correlation between HDL-C and TG. To test this, we changed plasma TG by varying LPL-activity, VLDL-PR or VLDL size and studied the resulting relationships between plasma TG and HDL-C.

Following expectations, decreasing LPL-activity led to a negative correlation between TG and HDL-C (Fig. 4A). In contrast, when VLDL-PR and VLDL-size are increased in physiological ranges, we unexpectedly observed a positive relation between plasma TG and HDL-C (Fig. 4B, Fig. 4C). Thus the model results oppose the idea that increased VLDL-PR causes lower HDL-C.

**Fig 4.**
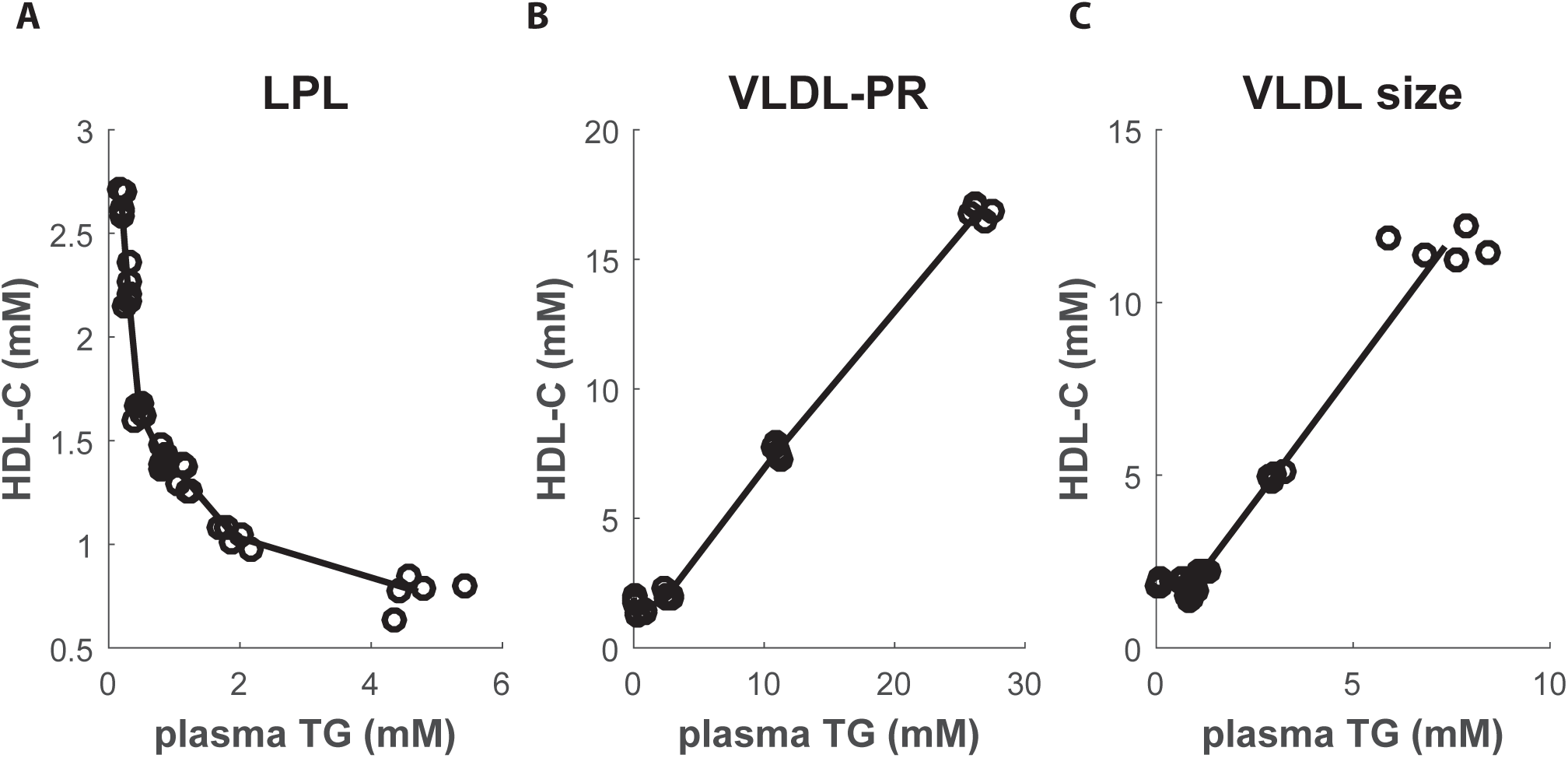
Effect of perturbing plasma TG on relation between TG and HDL-C.

The relation between HDL-C and plasma TG upon perturbing the system by varying LPL-activity, VLDL-PR and VLDL size respectively. Clearly, decreasing LPL leads to a negative correlation while increasing VLDL-PR and VLDL size leads to a positive correlation for the larger part of the parameter range.

To study this discrepancy, we studied the effect of VLDL-PR and VLDL-size on the apoA-I FCR in our model. We then found that VLDL-PR correlated positively with apoA-I FCR between [0 – 1] of normal VLDL-PR, while correlating negatively between [1 – 10] times of normal VLDL-PR (Fig. 5; panel A). Of note, this includes the VLDL-PR range observed in tracer-kinetic study in obese patients [0.75 - 3.0] [51–53]. Similarly, when modulating the size of VLDL through increasing the TG-content of VLDL produced, the correlation between VLDL-size (i.e. TG content) and apoA-I FCR turns from positive to negative beyond normal-sized VLDL (Fig. 5; panel B).

**Fig 5.**
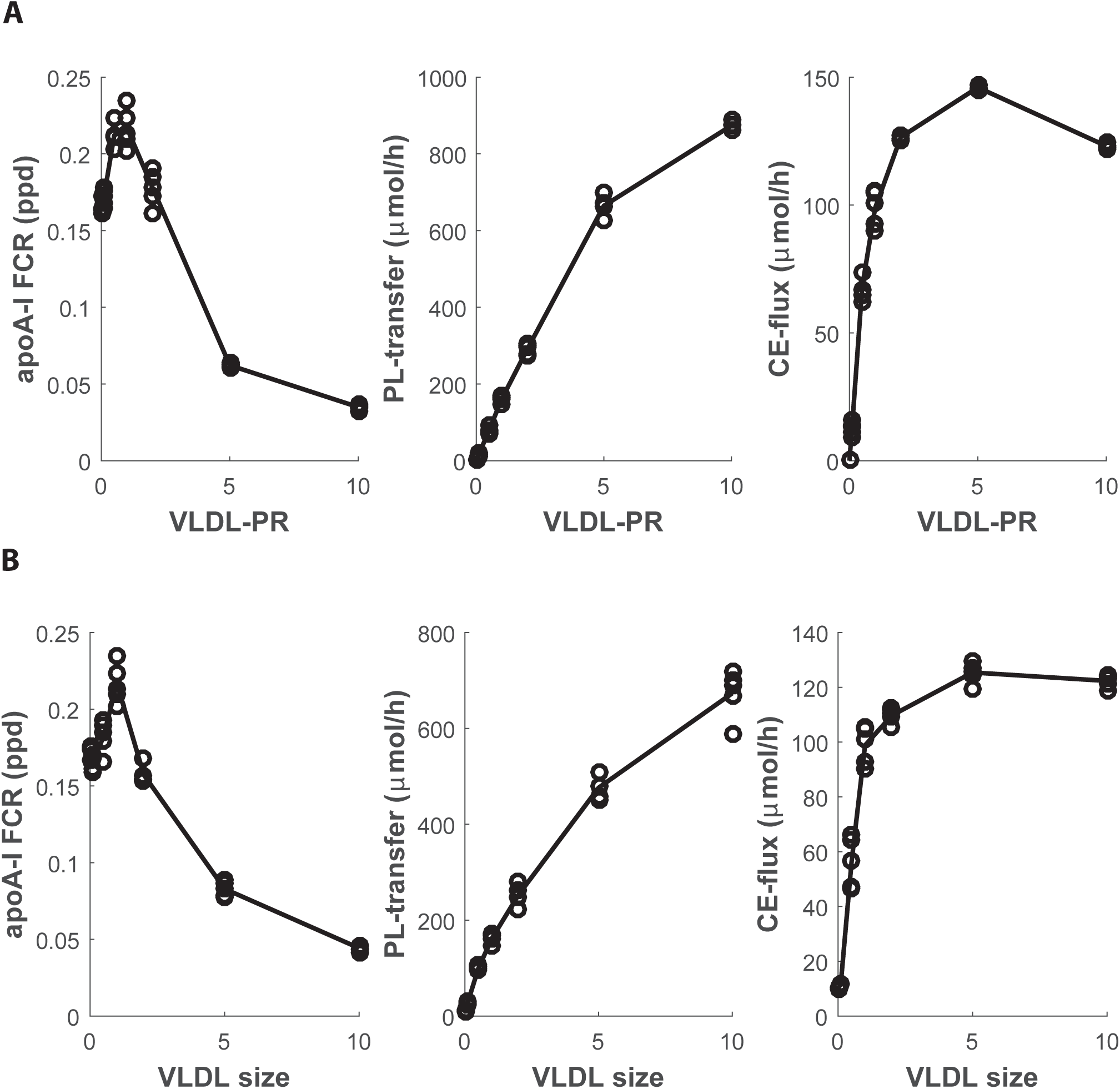
Effect of perturbing plasma TG on lipid fluxes between VLDL and HDL.

Effect of perturbing plasma TG through modulating VLDL-PR (A) and VLDL size (B) on apoA-I FCR respectively, PL-transfer rate, and CE-flux, respectively. Note how there is a maximum in apoA-I FCR for both VLDL-PR and VLDL-size.

We hypothesized that at higher ranges of VLDL-PR, the model reacts with a greater increase in LCAT-activity in response to increased PL-flux from VLDL to HDL compared to the increase in the CETP-mediated CE-flux from HDL to VLDL, leading to positive associations between HDL and TG. Conversely, we hypothesized that CE-flux from HDL to VLDL increases faster than the increase in LCAT-activity through additional supply of PL in the low range of VLDL-PR. To address these ideas, we inspected the PL-transfer and CE-flux over the studied parameter range and observed that, consistent with this hypothesis, while CE-flux flattens at higher rates of VLDL-PR, the increase in LCAT-activity in response to increased PL-flux remains strong in this range (Fig. 6A, 6B).

**Fig 6.**
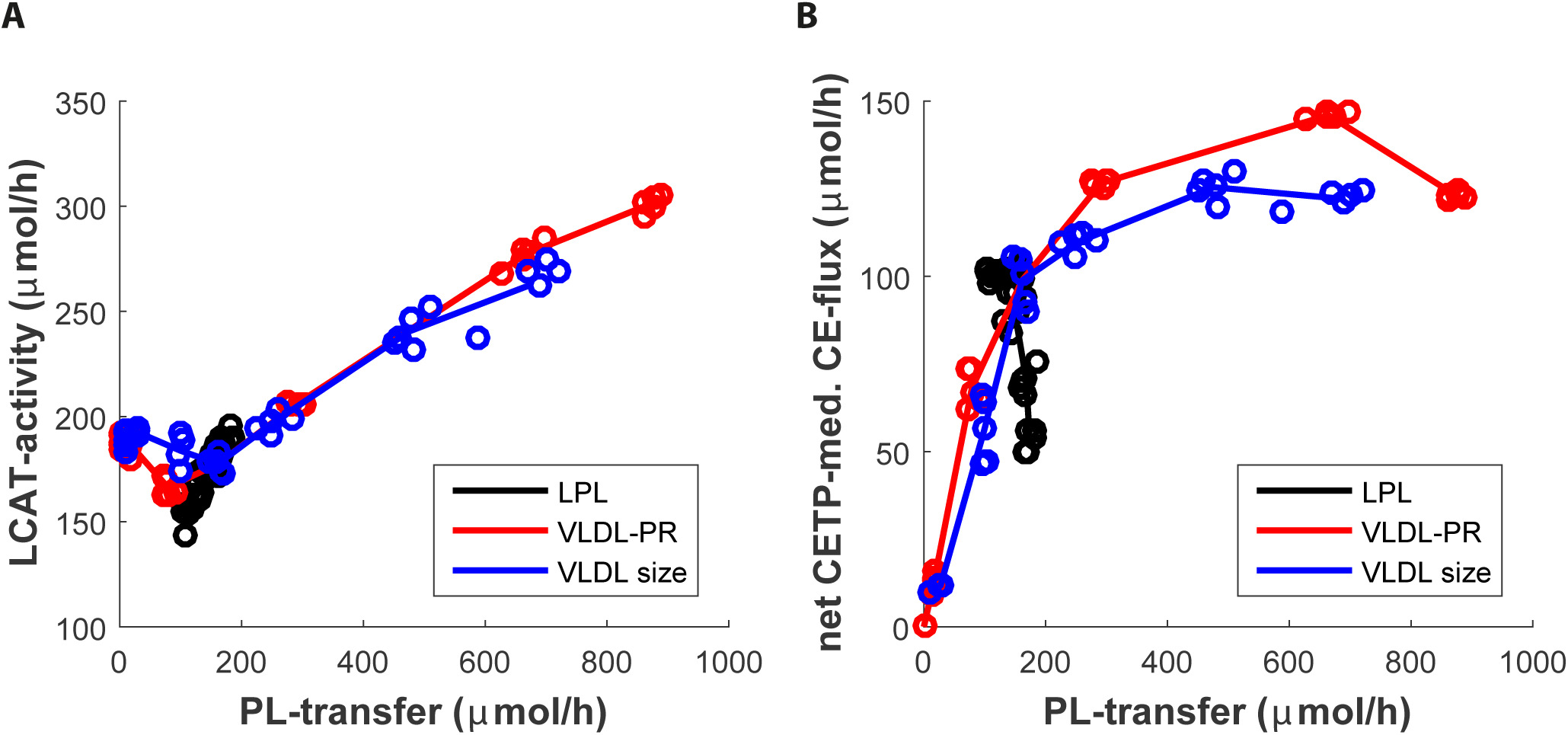
Effect of PL-transfer from VLDL to HDL on LCAT-activity and CE-exchange.

The relation between PL-transfer and CE-A-production, i.e. LCAT-activity (A), and between PL-transfer and CE-flux (CETP-mediated) from HDL to VLDL (B) when perturbing LPL-activity, VLDL-PR and VLDL size respectively. Note how there is a tight association between the phospholipid flux and LCAT-activity, and that at higher VLDL-PRs, PL-flux still increases while CE-flux does not.

The conclusion of this part of our study is that for VLDL-PR, not the CE-flux, but the PL-flux becomes dominating in determining the TG-HDL relationship.

### Positive HDL-TG correlation with increased VLDL-PR is insensitive to CETP

As indicated, the modeling result that increased VLDL-PR and VLDL-TG content do not lead to decreased plasma HDL-C is not in agreement with findings from tracer-kinetic studies [23,51,57]. Given this discrepancy, we re-evaluated how CETP-function was modeled, which after all, is one of the central players in the connection between HDL and TG metabolism. We first explored a wide range of Vmaxes for the CETP-algorithm. We then found that for all parameter combinations the positive association between HDL-C and plasma TG remained (Suppl. Fig. S3). Next, we considered whether we may have chosen a wrong algorithm for CETP. We compared the results of the current algorithm used, against results with two other algorithms we had considered for CETP in previous iterations of the model. We found that for both other CETP-algorithms, there was a negative association between apoB production and the apoA-I FCR, resulting in a positive association between HDL-C and plasma TG (Fig. 7). Taken together, none of the CETP-algorithms tested was able to negate the positive relation between TG and HDL-C at increasing VLDL-PR. This exercise therefore suggests that this modeling result is relatively insensitive to the CETP-algorithm used.

**Fig 7.**
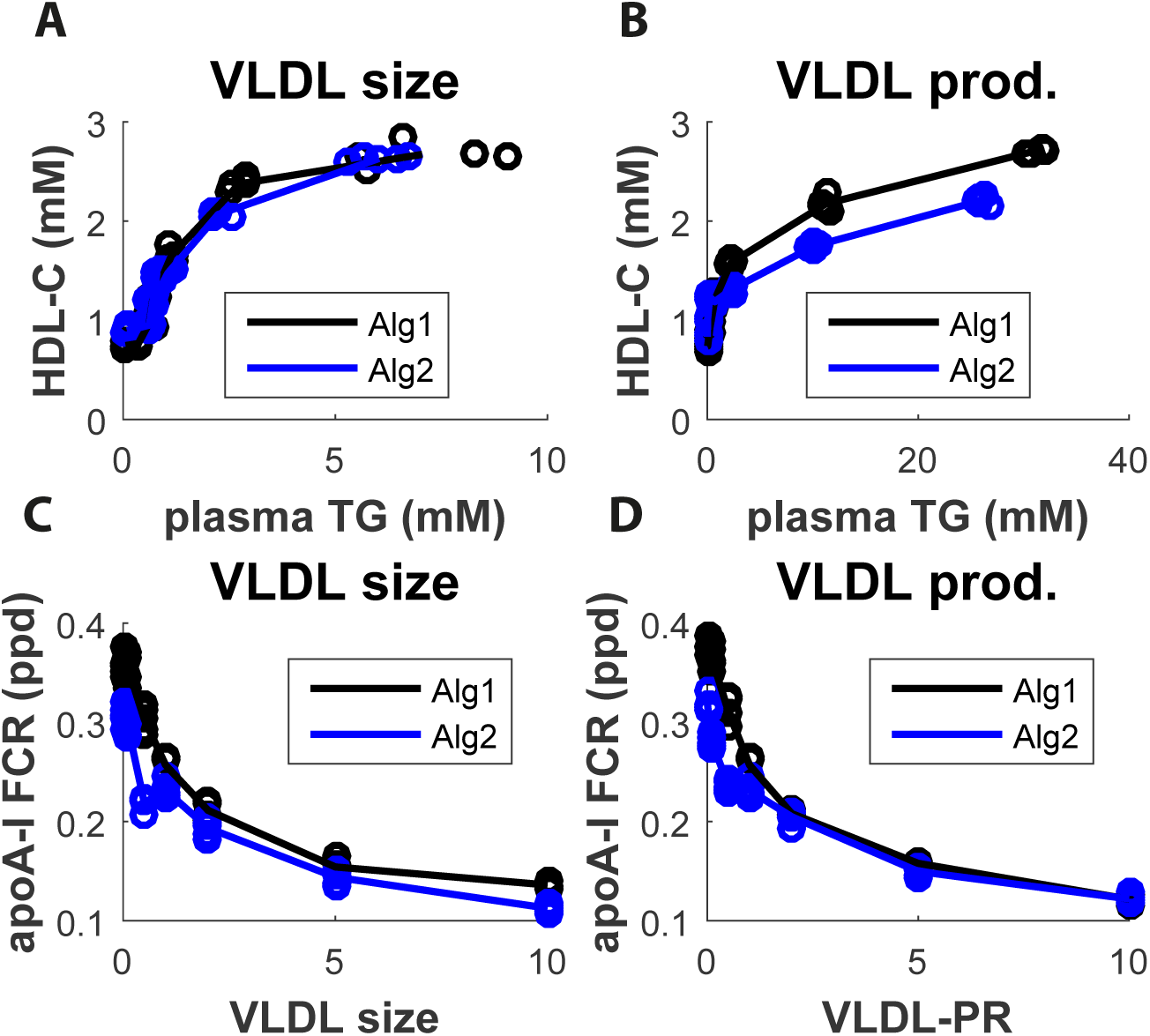
Effect of alternative CETP-algorithms on relation between plasma TG and HDL-C.

The relation between plasma TG and HDL-C, and the effect on the apoA-I FCR, when varying the size of VLDL produced and VLDL PR for two alternative algorithms for CETP (Alg1, and Alg2). VLDL size = 1, and VLDL-PR = 1, reflects production of normal (default) size VLDL and a normal VLDL-PR respectively. Note how the use of the two alternative algorithms for CETP, as the CETP-algorithm chosen for our model also leads to a positive correlation between HDL-C and plasma TG, and a decreasing apoA-I FCR when increasing VLDL production or size. Thus, the CETP-algorithm used has no effect on this modeling result.

### Addressing discrepancy with tracer-kinetic models and parameter ranges

Inspection of the modeling outcome suggested that our results could possibly be fitted with the findings in tracer-kinetic studies when the amount of PL available to apoA-I (or the HDL pool) would be less at any given rate of apoB-production. To test whether decreased availability of PL would indeed reverse the identified positive relation between TG and HDL-C, we repeated simulations with 10% and 30% instead of 65% of PL transfer from the excess surface of apoB-lipoproteins to HDL. Interestingly, this led to positive correlations between both VLDL size or VLDL-PR with the apoA-I FCR for a greater range of parameter values. Conversely, when the phospholipid transfer is increased to 90% (black line), this range is markedly reduced or even minimized (Fig. 8A, Fig. 8B). More importantly, decreasing the PL-transfer negates the positive correlation between plasma TG and HDL-C when VLDL-size is varied (Fig. 8C). Moreover, the correlation between plasma TG and HDL-C induced through modulation of VLDL-PR then turns from positive to negative (Fig. 8D). Finally, repeating the simulations for loss-of-function mutations in major lipid genes (*LCAT, ABCA1, APOA1, SCARB1, CETP, LDLR*) with a PL transfer of 10% instead of 65%, yield results that compare equally well with patient data (Fig. S4). Taken together, these findings suggest that the majority of excess surface lipids from VLDL may not be cleared through HDL.

**Fig 8.**
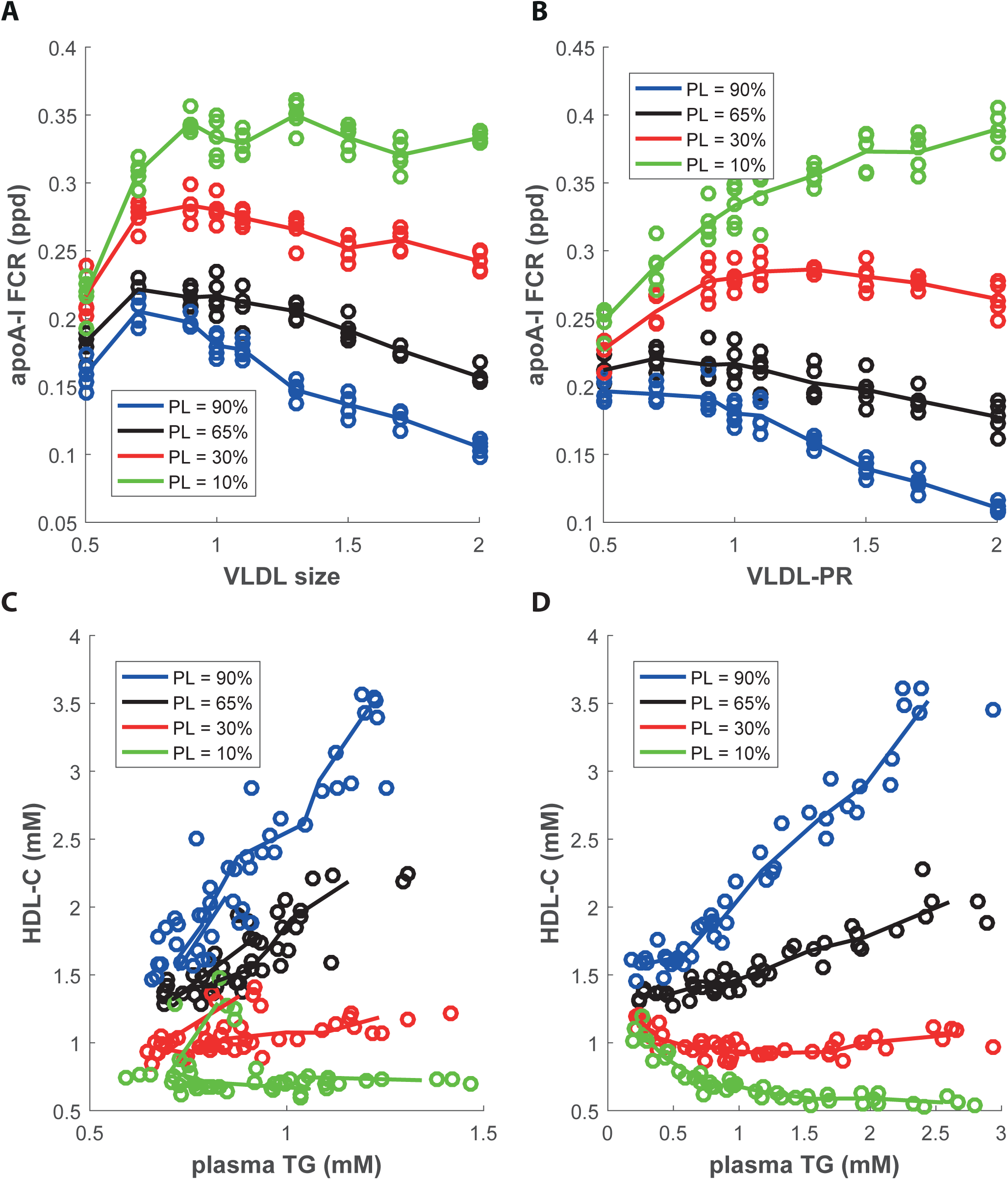
Effect of changing percentage of VLDL excess surface lipids transferred to HDL.

The effect of changing VLDL size and VLDL-PR on both the apoA-I FCR (A, B), and the TG-HDL-C relationship (C, D) when assuming 90%, 65%, 30% and 10% availability of PL to HDL respectively. Note how decreasing, or increasing the PL-flux to HDL shifts the relation between VLDL-PR and apoA-I FCR, as well as the relation between VLDL size and apoA-I FCR to the right and left respectively. Similarly, the positive correlation between plasma TG and HDL-C turns neutral for increasing VLDL size and turns negative for increasing VLDL-PR.

### Is there an HDL independent pathway for the clearance of excess surface lipid from VLDL?

Our findings thus suggest that a majority of surface lipids from VLDL shed through lipolysis may not be cleared through HDL. This raises the question through what route the majority of the excess surface lipids, if not through HDL, should then take. To our knowledge, research on alternative pathways for PL transfer from VLDL or chylomicrons to other particles than HDL has not been published. It is in this regard interesting to consider what happens with the excess surface lipids in familial HDL deficiencies, where ‘clearance of PL’ through HDL is not possible:

Specifically, complete loss of LCAT activity (familial LCAT deficiency; FLD) causes the accumulation of PL in plasma in the form of lipoprotein X (LpX) that is assumed to originate from the pinching off of surface lipids from VLDL [58]. LpX are lamellar aggregates consisting of mainly PL, FC and some albumin [59]. Remarkably, patients suffering from partial LCAT deficiency (fish-eye disease; FED) also suffer from HDL deficiency but they do not present with LpX [59,60]. Likewise, patients with a complete loss of ABCA1 activity (Tangier Disease) or apoA-I deficiency, again both characterized by HDL deficiency, do not present with LpX either [61–63]. The fact that LpX is present in only FLD while absent in three other cases of HDL-deficiency, leaves room for the idea the substrates that accumulate into LpX, are unlikely to be cleared through HDL, but may be cleared through an alternative pathway.

Interestingly, LpX are also observed in patients with cholestasis, where bile (containing PL and FC) may enter the plasma compartment [64]. So far, however, it is unclear what causes the accumulation of LpX in either FLD or cholestasis.

It is tempting to speculate that macrophages may play an important role in the clearance of these excess surface lipids, and that the fact that LpX is not observed in Tangiers disease results from enhanced uptake of LpX in this specific condition. There are indications, however, that macrophage lipid accumulation in Tangier disease rather relates to the inability to metabolize certain lipid species, due to the absence of ABCA1 affecting lysosomal function [65].

Activity of HL and EL on apoB-proteins is not likely to be the major pathway either, since activities of these enzymes are normal in FLD, yet do not prevent the formation of LpX. It is in this regard more likely that beta-LCAT activity, i.e. residual activity of LCAT on only LDL as described for FED, is sufficient to prevent the formation of LpX [66]. Together, these observations argue for a system in which LCAT-activity is necessary for the removal of surface lipids from VLDL that cannot be accounted for by a flux through HDL.

## Concluding remarks

Making use of established knowledge of human lipoprotein metabolism, including tracer-kinetic data on HDL and VLDL-metabolism, we have constructed an agent-based model that can adequately simulate plasma lipid and lipoprotein measurements in heterozygotes and homozygotes of loss-of-function mutations in *LCAT, ABCA1, APOA1, SRB1, CETP* and *LDLR*. The model was built to specifically study the etiology of the inverse relation between plasma TG levels and HDL-C. This intriguing relationship is generally seen in observational as well as intervention studies (CETPi, Niacin, LPL gene therapy) of lipoprotein metabolism and cardiovascular disease [67–70]. Remarkably, however, the mechanism underlying this relationship is unclear.

The general contention is that the majority of excess VLDL surface lipids during TG hydrolysis are transferred to HDL. Our simulations, however, show that even a conservative estimate of a 65% transfer of excess surface lipids to HDL would actually increase HDL-C concentration at physiological ranges of increased plasma TG. In fact, to simulate a negative association between HDL-C and plasma TG, PL-transfer had to be limited from 65% to 10% to 30%. This result point at an alternative pathway that is responsible for the bulk of excess PL removal.

In this regard, it is interesting that the generation of LpX, causing renal failure and early death in patients with FLD, is assumed to be the result of impaired clearance of excess PL. The question is, why do these lipid agglomerates not occur in other familial HDL-C deficiencies such as apoA-I deficiency, Tangier disease or FED (partial LCAT deficiency)? This interesting question has to our knowledge hardly received attention.

Our study shows that agent based modeling is a useful tool to unravel the complex interplay between lipoproteins in human plasma. Furthermore, our findings suggest that an alternative HDL-independent pathway for the disposal of PL following TG hydrolysis of VLDL may be of clinical and pharmaceutical relevance.

## Acknowledgements

We thank D.J. Reijngoud for helpful discussions. This work was supported by grants from the European Union grant FP7-HEALTH n°305707; acronym RESOLVE (YP, JAK and AKG) and FP7-603091; Acronym TransCard as well as the Netherlands CardioVascular Research Initiative: “the Dutch Heart Foundation, Dutch Federation of University Medical Centers, the Netherlands Organization for Health Research and Development and the Royal Netherlands Academy of Sciences” (CVON2017-2020; Acronym Genius2 to JAK). JAK is Established Investigator of the Netherlands Heart Foundation (2015T068).

## Supporting Information

**Suppl. File S1 – Model Description**

**Suppl. File S2 – Simulation Files**

**Suppl. Fig S1 – Sensitivity Analysis plasma TG**

**Suppl. Fig S2 – Sensitivity Analysis HDL-C**

**Suppl. Fig S3 – Effect of Changing Vmaxes CETP-algorithm on HDL-TG relation Suppl. Fig S4 – Monogenetic Disorder Simulations with PL-transfer of 10%**

